## Supplementary Information for "A pH-dependent cluster of charges in a conserved cryptic pocket on flaviviral envelopes"

**Supplementary Figures**

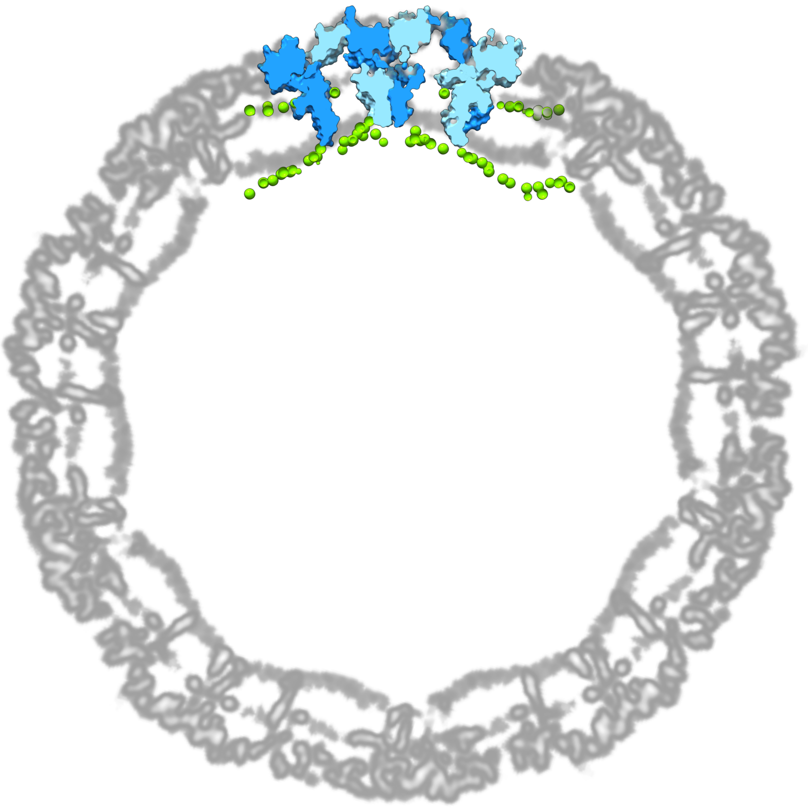

**Figure S1. An atomistic model of a DENV2 raft and its curved membrane overlaid with a cryo-EM map of a DENV2 envelope** (PDB: 3J27; EMDB: EMD-5520). The EM map is depicted as a single plane spanning the centre of the viral particle. For clarity, the raft model is visualised as a thin slice overlaid with the cryo-EM map. Raft chains are shown in alternating light and dark blue surfaces, while membrane phosphorus atoms are shown in green spheres.

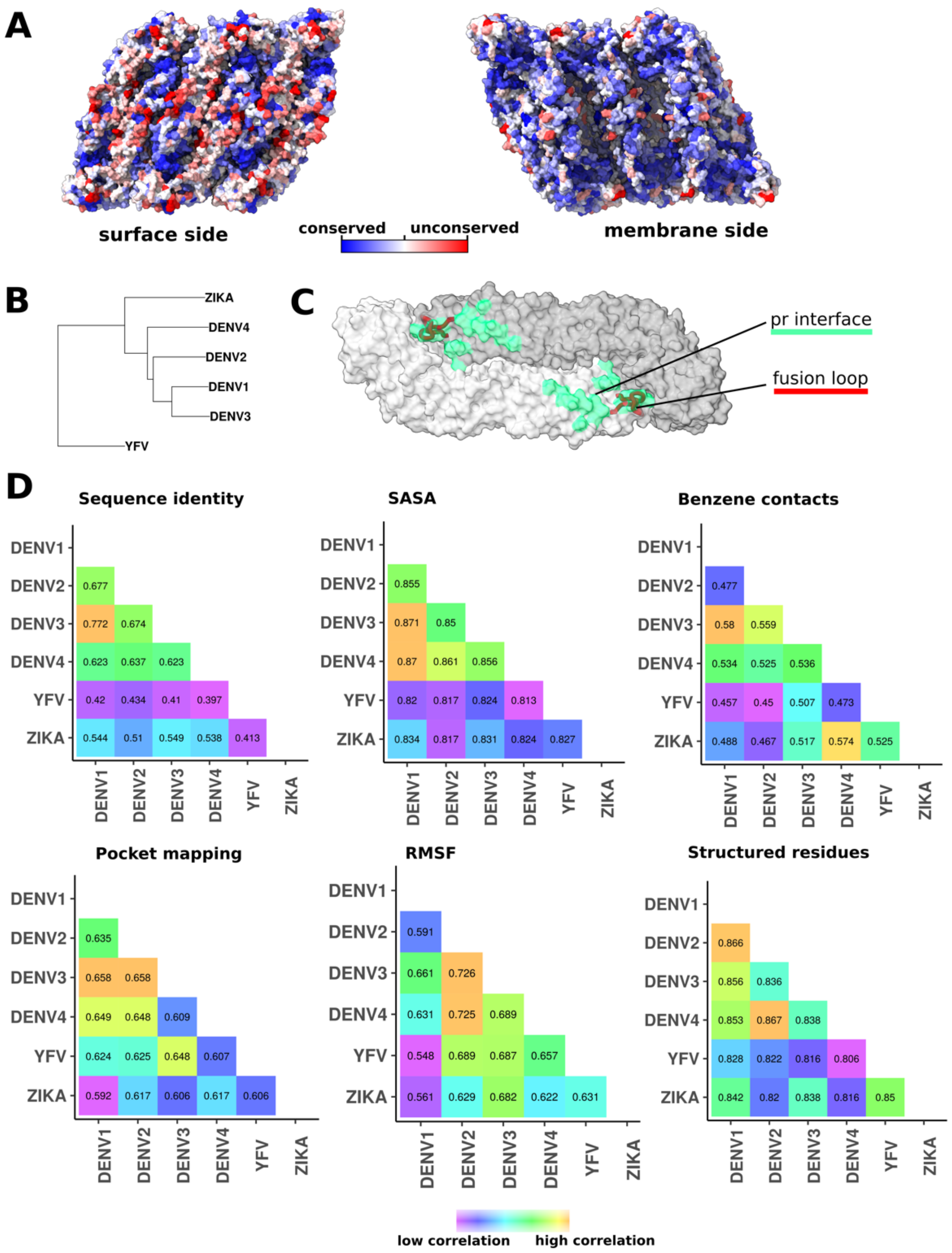


**Figure S2. Residue conservation and phylogenetic relationships of flaviviruses correlated with simulation properties. A)** Raft residue conservation as shown on the raft surface. The surface side is turned towards the exterior of the viral particle, while the membrane side is interacting with the lipid bilayer and is for the most part hidden from the host immune system. **B)** Phylogenetic relationship of six flaviviruses presented by an unrooted neighbour joining (NJ) tree. DENV1 and DENV3 are most closely related viral serotypes, while YFV is comparatively distant in evolutionary terms to the DENV group. **C)** The E protein dimer showing the pr interface (green surface) and the fusion loop (red ribbon; residues 98-111). The dimer is shown in a surface representation with individual chains coloured in shades of grey. **D)** Correlation analyses of properties associated with the selected viral serotypes. Sequence identity matrix shows percentage identity of E and M protein sequences in a pairwise fashion. This matrix reflects evolutionary relationships between viruses shown in panel B. Other matrices display pairwise correlations of viral properties obtained from MD simulations. The values of SASA, RMSF, and number of structured residues were derived from water-only simulations, while benzene contacts and pocket densities were taken from benzene simulations. All correlations were calculated using the Spearman's rank correlation coefficient method.

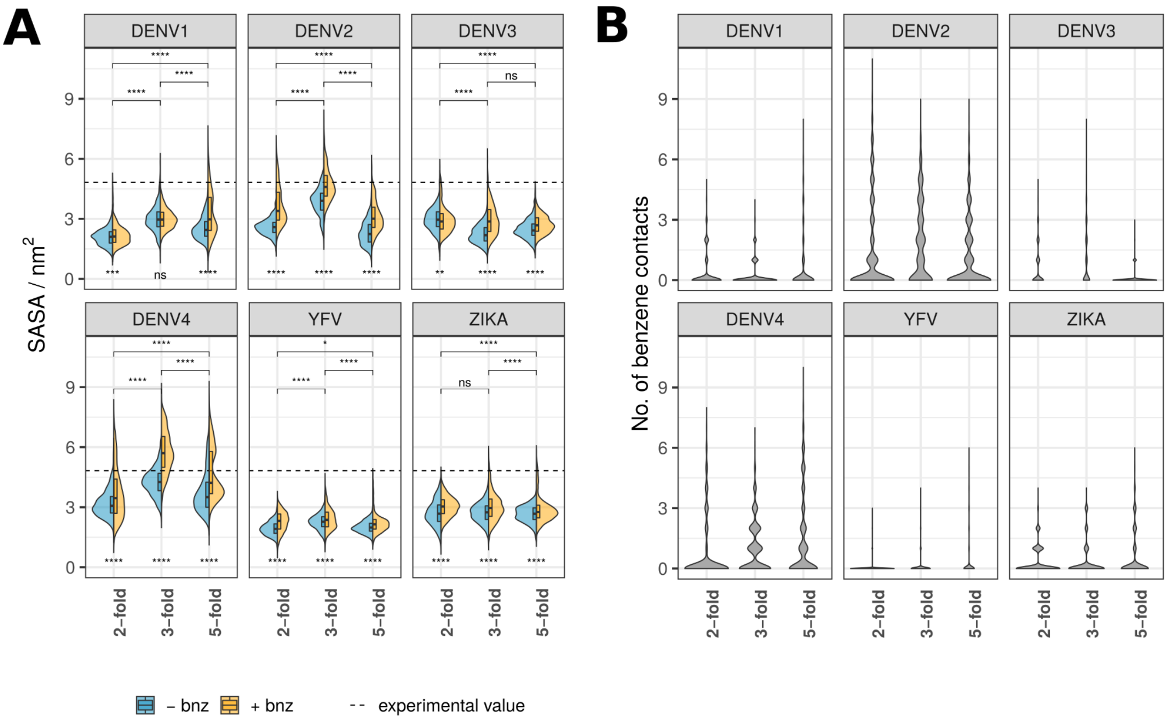


**Figure S3. β-OG pocket properties of SASA and benzene contacts shown for individual viral strains and chain types. A)** SASA calculated for β-OG pocket residues shown in split violin plots, where the blue area represents pocket SASA in water-only simulations, while the yellow surface represents simulations with benzene molecules present in the solvent. Experimental SASA value corresponds to the crystal structure pockets averaged across two chains. The division of pockets based on chain positions on the raft follows the labelling convention described in Figure 1A. Scores underneath the violin plots indicate the significance of SASA difference for water-only and benzene simulations. Pairwise scores above the plots describe a significant difference between chain types (across both solvent types). Significance is calculated using a Wilcoxon signed-rank test (ns: $p > 0.5$; *: $p \leq 0.5$; **: $p \leq 0.01$; ***: $p \leq 0.001$; ****: $p \leq 0.0001$). **B)** The number of benzene contacts occupying the β-OG pocket at any given point throughout the course of simulations. The values are shown for individual viruses and chain types.

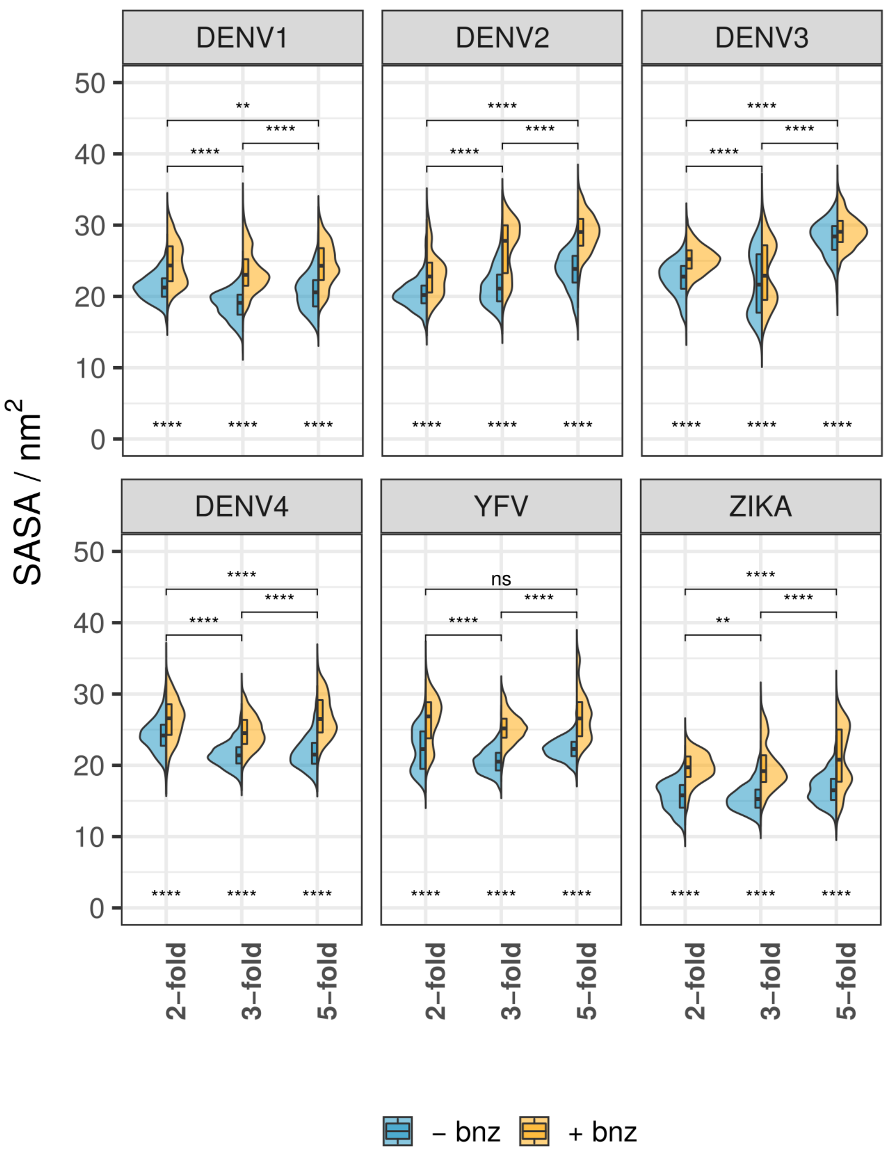


**Figure S4. α pocket SASA values across different viral strains and solvent compositions.** Grouping categories and the methods for calculating statistical significance are described in Figure S2A.

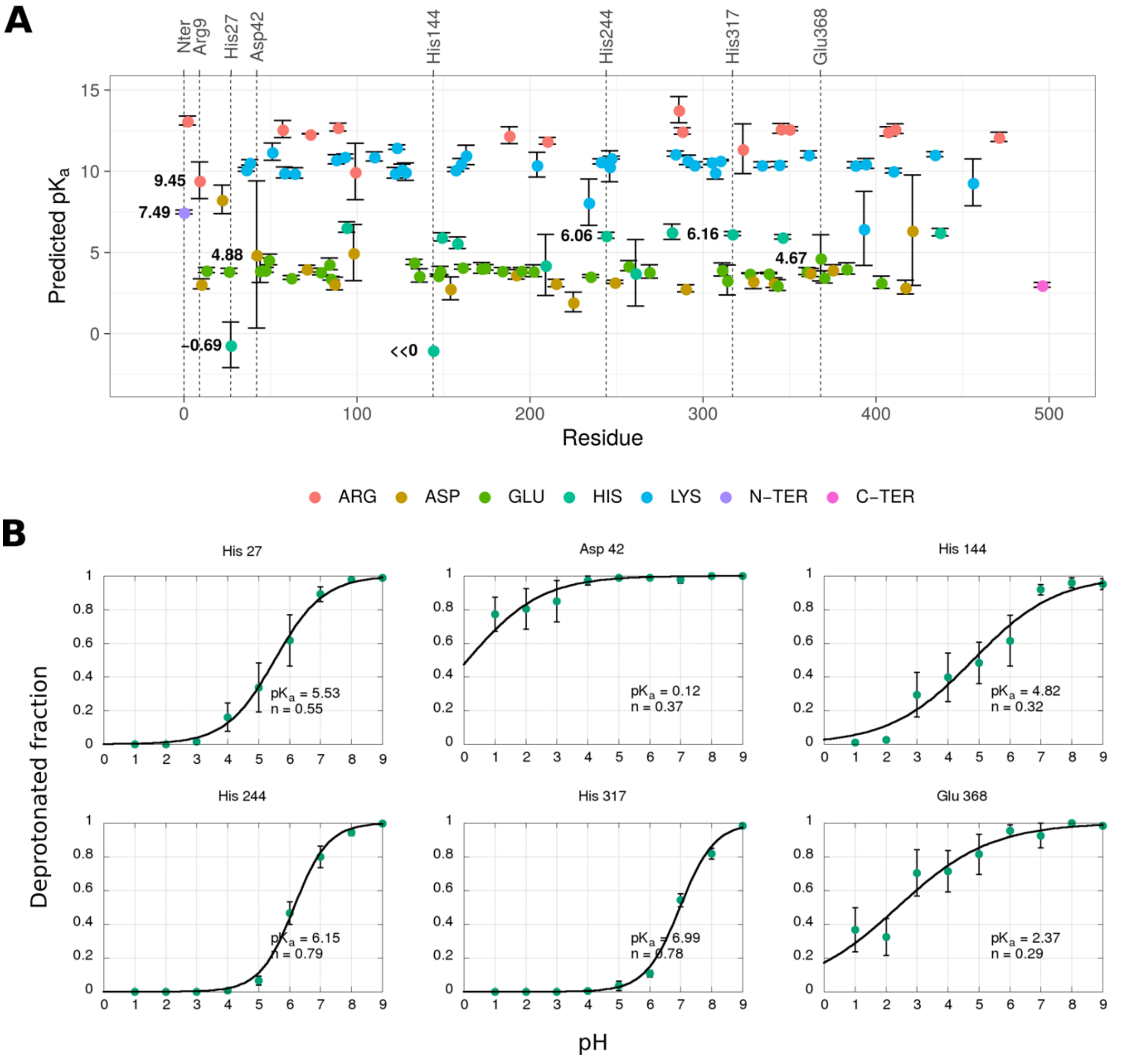


**Figure S5. Prediction of p*K*a values for selected E protein residues using static structures and MD simulations. A)** A p*K*a prediction for all titratable residues of the E protein calculated by the FD/DH method. The values are averaged across three chains in the asymmetric unit and standard errors are shown as whiskers. **B)** Titration of selected residues derived from constant-pH simulations of the DENV2 sE dimer. The values are averaged across both chains and all repeats, with the standard errors shown as whiskers. The curves are fitted according to the Hill equation, where *n* is a Hill coefficient describing cooperativity between proton binding sites (n < 1, negative cooperative binding; n = 1, independent binding; n > 1, positive cooperative binding).

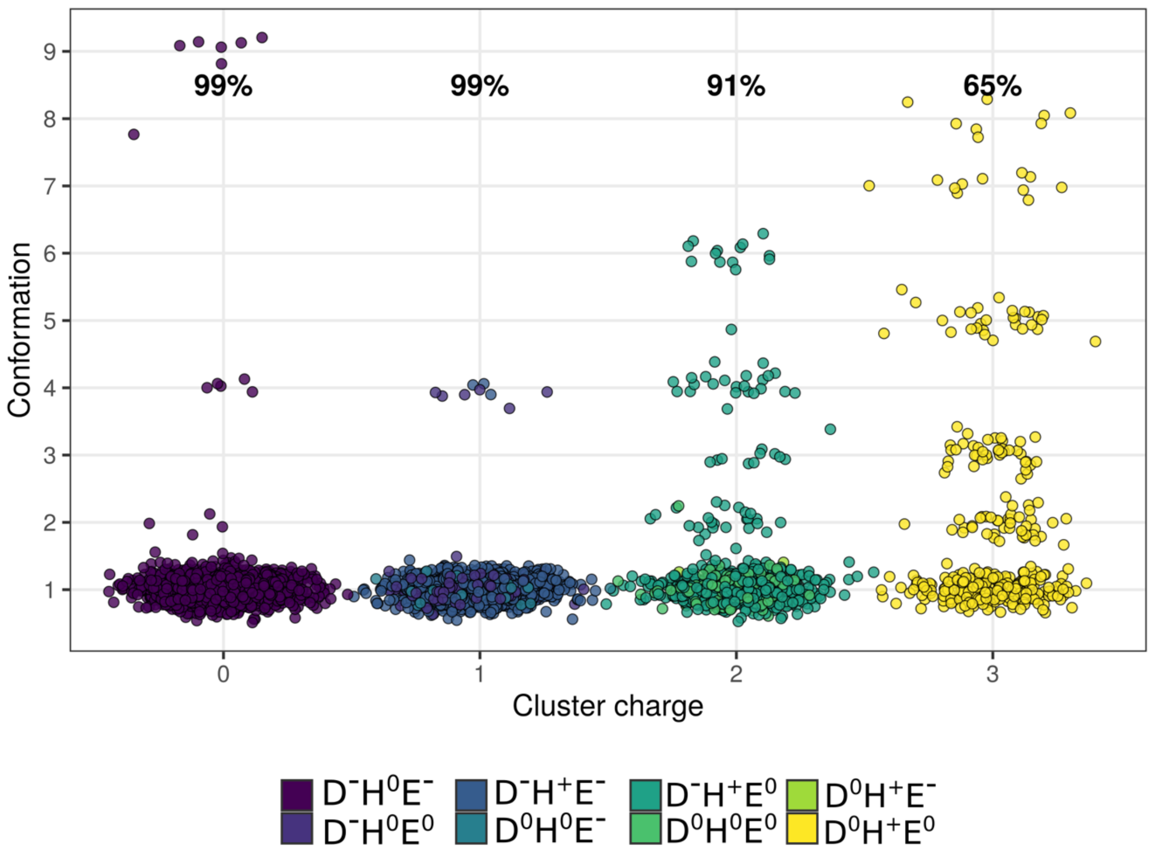


**Figure S6. Occurrence of conformations at different charge states.** Conformation 1 is predominant at all charge states, whereas other conformations are enriched at higher charges (+2 and +3) occurring at low pH. The labelled percentages describe the population proportion belonging to conformation 1.

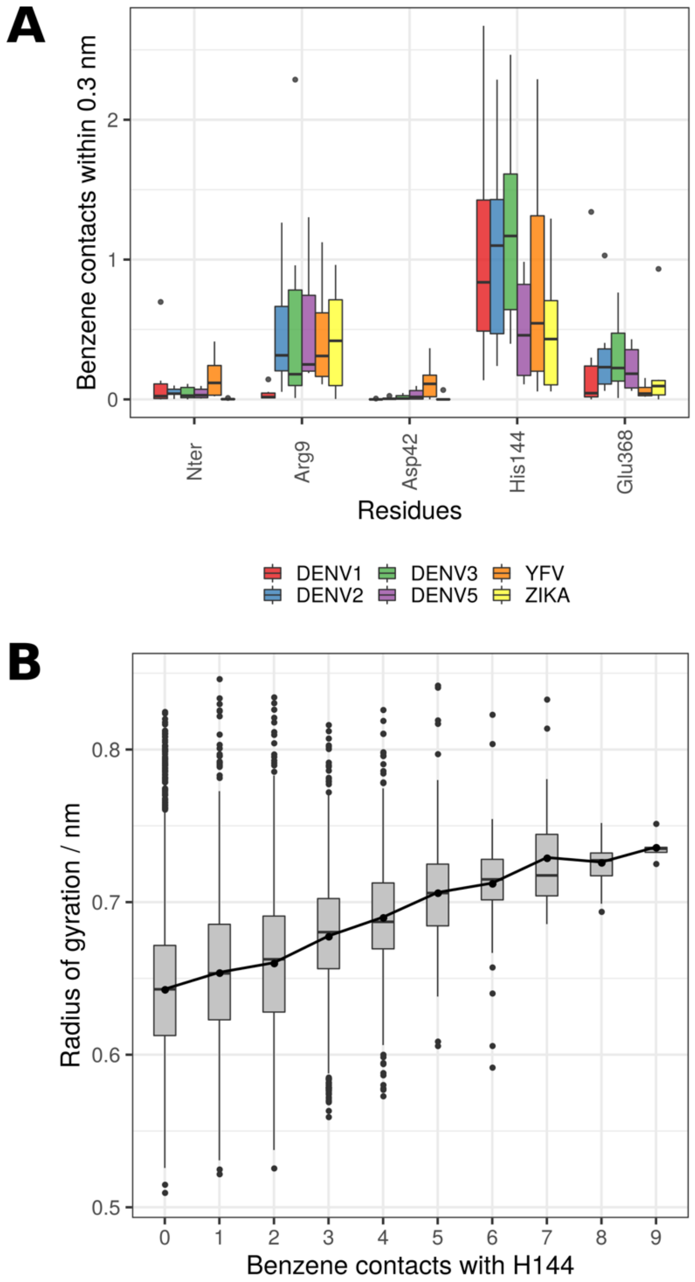


**Figure S7. Benzene contacts and its effects on the charge cluster. A)** Number of benzene contacts for each residue within the cluster of charges (Nter-Arg9-Asp42-His144-Glu368) and across examined virus species. **B)** The radius of gyration (measuring cluster disruption) in relation to the number of benzene molecules interacting with His144 across all examined flaviviruses. The increased number of benzene molecules within the α pocket leads to a greater disorder within the cluster. A point-to-point line connects the mean values of all groups (whereas boxplots highlight the median values of groups).

### Supplementary Videos

**Supplementary Video 1. The top-down view of the DIII dissociation in DENV2 sE dimer under low pH conditions.** The trajectory corresponds to the snapshot shown in yellow ribbons in Figure 6C. The domains are coloured as in Figure 1C, and the residues within the cluster of charges are shown in licorice. Transparent surfaces represent the starting sE dimer conformation.

**Supplementary Video 2. The side view of the DIII tilt in DENV2 sE dimer under low pH conditions.** The trajectory corresponds to the snapshot shown in green ribbons in Figure 6C. The domains are coloured as in Figure 1C , with the charge cluster residues shown in licorice. Transparent surfaces represent the starting sE dimer conformation.
